# A note on the analysis of two-stage task results: how changes in task structure affect what model-free and model-based strategies predict about the effects of reward and transition on the stay probability

**DOI:** 10.1101/187856

**Authors:** Carolina Feher da Silva, Todd A. Hare

**Author notes:** Corresponding author, Laboratory for Social and Neural Systems Research, Department of Economics, University of Zurich, Zurich, Switzerland.

## Abstract

Many studies that aim to detect model-free and model-based influences on behavior employ two-stage behavioral tasks of the type pioneered by Daw and colleagues in 2011. Such studies commonly modify existing two-stage decision paradigms in order to better address a given hypothesis, which is an important means of scientific progress. It is, however, critical to fully appreciate the impact of any modified or novel experimental design features on the expected results. Here, we use two concrete examples to demonstrate that relatively small changes in the two-stage task design can substantially change the pattern of actions taken by model-free and model-based agents as a function of the reward outcomes and transitions on previous trials. In the first, we show that, under specific conditions, purely model-free agents will produce the reward by transition interactions typically thought to characterize model-based behavior on a two-stage task. The second example shows that model-based agents’ behavior is driven by a main effect of transition-type in addition to the canonical reward by transition interaction whenever the reward probabilities of the final states do not sum to one. Together, these examples emphasize the task-dependence of model-free and model-based behavior and highlight the benefits of using computer simulations to determine what pattern of results to expect from both model-free and model-based agents performing a given two-stage decision task in order to design choice paradigms and analysis strategies best suited to the current question.

## 1 Introduction

The brain contains multiple systems that interact to generate decisions, among them model-free systems, which reinforce rewarded actions and create habits, and model-based systems, which build a model of the environment to plan toward goals. Model-free and model-based influences on behavior can be dissociated by multi-stage behavioral tasks. In such tasks, agents predict different state-action-reward contingencies depending on whether or not they employ a model of the task, i.e., whether or not they know how the transitions between task states most often occur [1]. Since the original two-stage task was first proposed and reported by Daw et al. [1], it or one of its variations has been employed by many studies on decision making (e.g., [2, 3, 4, 5, 6, 7, 8, 9, 10, 11, 12, 13, 14, 15, 16, 17]).

In the original two-stage task [1], each trial takes the participant sequentially through two different environmental states, where they must make a choice (Fig 1). Typically, at the initial state, the participant makes a choice between two actions, which we will refer to as “left” or “right.” Each initial-state action has a certain probability of taking the participant to one of two final states, which will be called “pink” and “blue.” Importantly, each initial-state action has a higher probability (for example, 0.7) of taking the participant to one of the final states, the “common” transition, and a lower probability (for example, 0.3) of taking the participant to the other final state, the “rare” transition. Let us assume that the left action commonly transitions to the pink state and the right action commonly transitions to the blue state. A participant should thus choose left if they want to maximize the probability of reaching the pink state and right if they want to maximize the probability of reaching the blue state. At the final state, the participant makes another choice between one or more actions (typically two), and each final-state action may or may not result in a reward with a certain probability. Typically, the probability of reward, or in some cases the reward magnitude, changes from trial to trial in order to promote continuous learning throughout the experiment.

**Fig 1:**
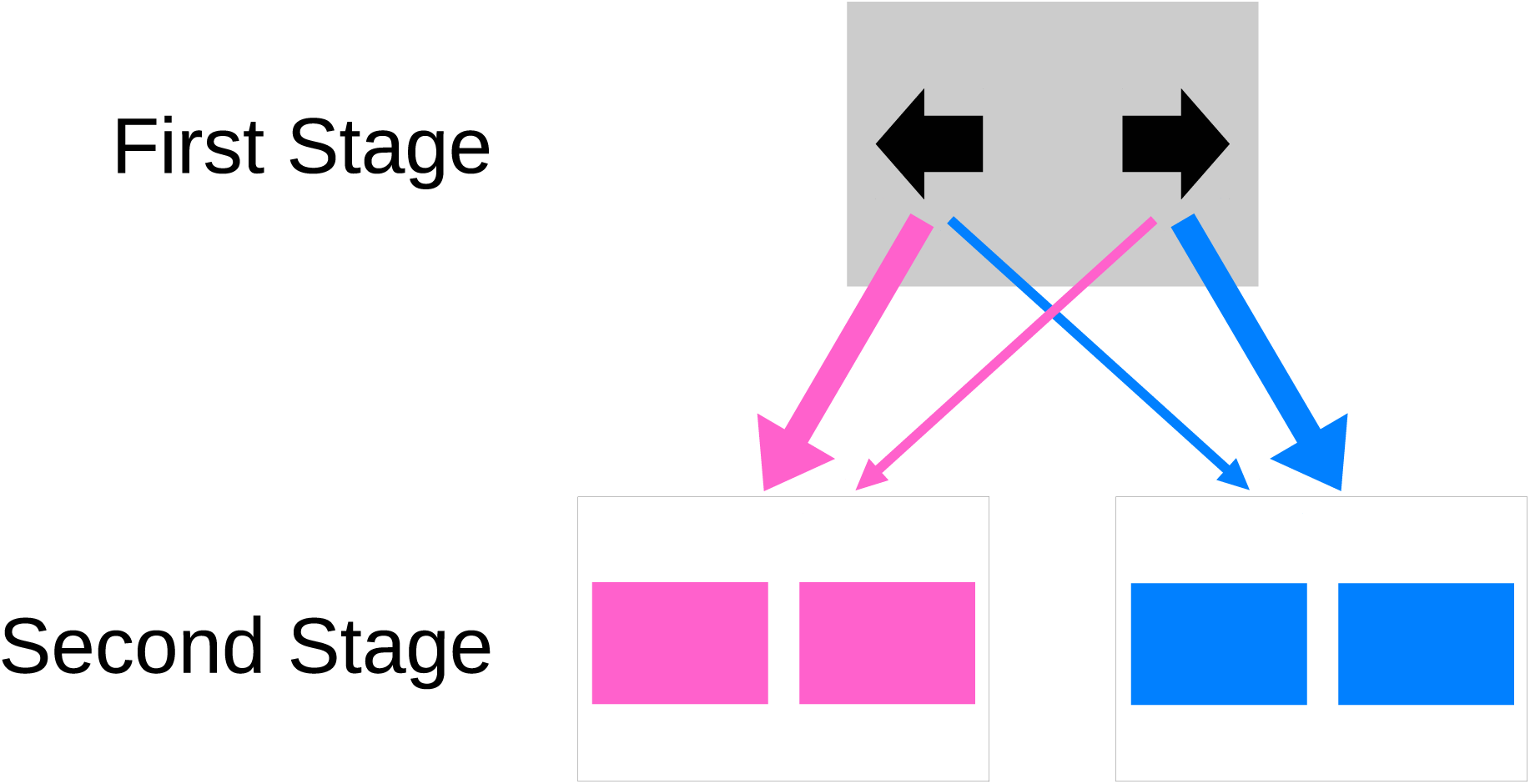
Scheme of a typical two-stage task. The thicker arrow indicates the common transition and the thinner arrow indicates the rare transition.

Daw et al. [1] proposed that, to analyze the results of this task, each initial-state choice is coded as 1 if it is a stay, that is, the participant has repeated their previous choice, or as 0 otherwise. Then, the participant’s stay probability is calculated depending on whether the previous trial was rewarded or not and whether the previous transition was common or rare. This analysis involves performing a logistic regression in which the stay probability is a function of two factors, reward and transition.

Applying this analysis to results obtained from simulated model-free or model-based agents produces a plot similar to that shown in Fig 2A. (Note that the exact stay probability values depend on the simulated agents’ parameters.) It is observed that for model-free agents, only reward affects the stay probability, and for model-based agents, only the interaction between reward and transition affects the stay probability. This difference allows us to distinguish between model-free and model-based choices.

**Fig 2:**
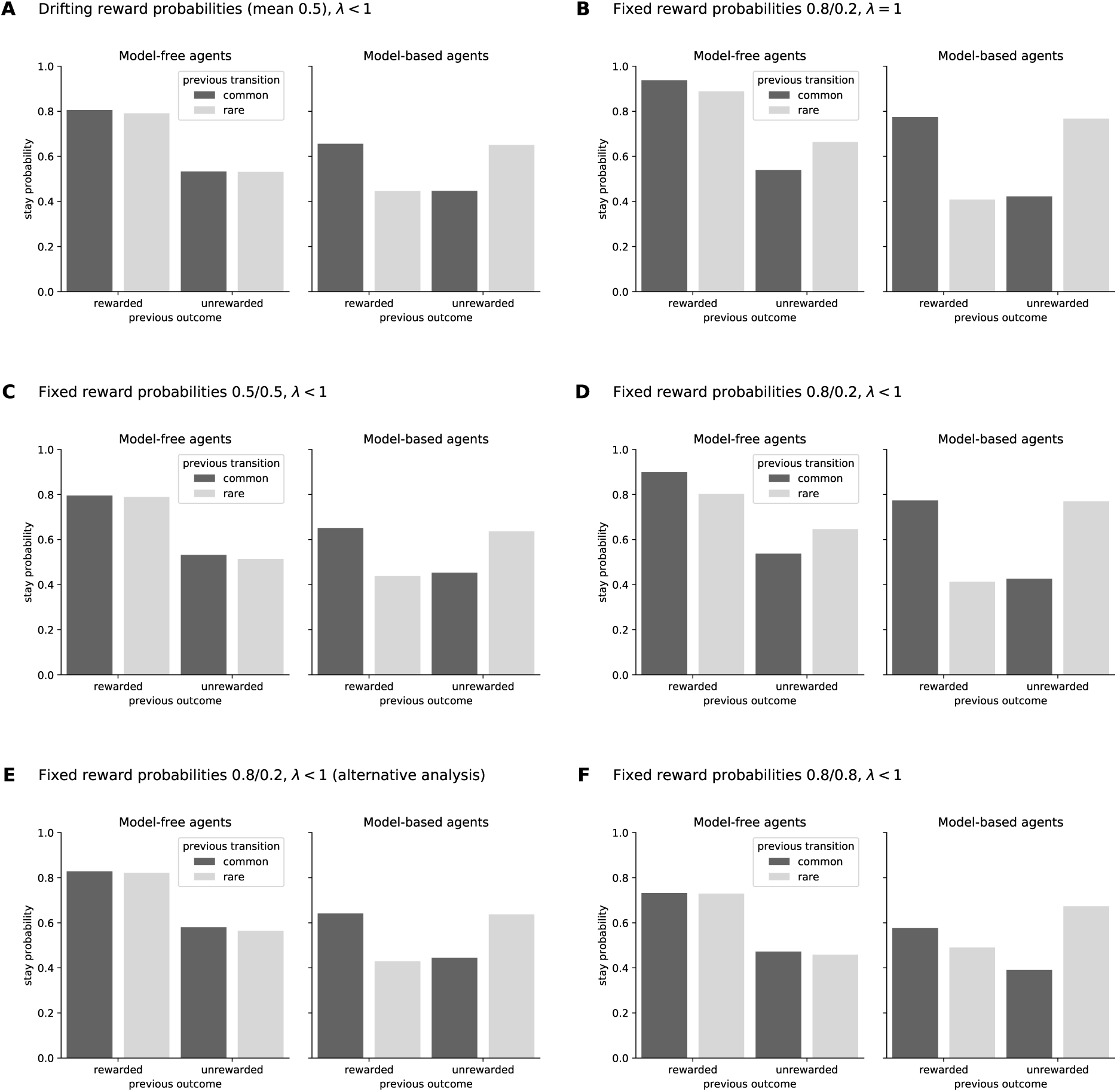
Results from the classical two-stage task as originally reported by Daw and colleagues (A) and variations (B–F), obtained by simulating model-free and model-based agents. In all panels, the behavior of simulated model-free agents are shown in the left bar-plots and model-based agents on the right. The y-axis shows the probability of staying with (i.e. repeating) the same action made on the previous trial. The x-axis separates the data as a function of previous outcome (rewarded, unrewarded) and transition (common = dark grey, rare = light grey). The data were analyzed by logistic regression, in which the stay probability was computed as a function of the previous outcome and transition, with the analysis in panel E) being modified to include additional regressors (see Section 2.1). The reward probabilities at each second stage and the agents’ eligibility trace (*λ*) are listed for each panel. **A)** The results from the classic two-stage task, as described by Daw et al. [1]. **B)** shows the pattern of stay probabilities when the second stage rewards are fixed at 0.8 and 0.2. **C)** is identical to panel A, except that both 2nd-stage reward probabilities are fixed at 0.5 instead of drifting independently around a mean of 0.5. **D)** is identical to panel B, except that the agents’ eligibility traces are set to values ¡ 1 instead of equal to 1. **E)** plots the same data as B), but analyzed with the extended logisitic regression discussed in Section 2.1. Lastly, **F)** presents the results of the modified task discussed in Section 2.2 in which the 2nd-stage reward probabilities sum to a value greater than 1.

The choice patterns of model-free and model-based agents in Fig 2A are different because model-based reinforcement learning algorithms take into account the task structure and model-free algorithms do not, with the result that they make different predictions about which action agents will choose at the initial stage. Here, we use “agent” as a general term to refer to either a computer simulation or a human or non-human participant. The model-free SARSA(*λ* = 1) algorithm predicts that if an agent makes a certain initial-state choice in a trial, they are more likely to repeat it on the next trial if it was rewarded, whether the transition was common or rare. A model-based algorithm [1], however, predicts that the agent is more likely to repeat the previous choice if, in the previous trial, it was rewarded *and* the transition was common, or if it was unrewarded *and* the transition was rare. For example, suppose an agent chooses left, is taken to the blue state through the rare transition, and receives a reward. In this case, the model-free prediction is that the agent is more likely to choose left again in the next trial, while the model-based prediction is that the agent is instead more likely to switch and chose right. The model-based agent is predicted to choose to go right, instead of left, at the initial state because the right action maximizes the probability of reaching the blue state, where the agent received the reward on the previous trial.

One might assume that even if the two-stage task structure is slightly changed to suit a particular research goal, model-free-driven actions will remain unaffected by transition-types because the model-free algorithm predicts that rewarded actions are more likely to be repeated regardless of transition. Similarly, one might assume that model-based choices will not be affected by reward because reward effects are characteristic of model-free actions. However, the general danger of relying on untested assumptions is well-known, and our work here aims to highlight the particular dangers of assuming fixed relationships between reward, transition-types, and model-free or model-based processing in two-stage tasks. It has already been demonstrated that these assumptions do not hold for a simplified version of the two-step task, optimized for animal subjects [15]. Here, we demonstrate by means of computer simulation that even seemingly small changes in task design can change the resulting choice patterns for model-based and model-free agents. For example, depending on the task details, it is possible that the stay probability of model-free agents is larger for common transitions than for rare transitions (i.e. that there is an interaction between reward and transition of the type thought to characterize model-based behavior). Below, we demonstrate two concrete examples of how slight changes in task design strongly affect the results of model-free and model-based agents in a logistic regression analysis. We also explain why these task features change the *expected* behavior of model-free and model-based agents and offer some further thoughts on how to analyze data from these modified tasks. Together, these examples emphasize the importance of simulating the behavior of model-free and model-based agents on any two-stage task, especially novel modifications, in order to determine what pattern of behavior to expect.

## 2 Results

### 2.1 Unequal reward probabilities make model-free agents indirectly sensitive to transition probabilities

Contrary to the assumptions of many researchers, it is not universally true that the stay probability of model-free agents is only affected by reward or that the stay probability of model-based agents is only affected by the interaction between reward and transition. Therefore, the stay probability plot will not necessary follow the “classic” pattern shown in Fig 2A; alterations in this pattern can stem from seemingly small and innocuous variations in the properties of the two-stage task.

The behavior of model-free agents is indirectly sensitive to the relative reward probabilities of the final states. If, for instance, we set the reward probabilities of the actions at the pink state to a fixed value of 0.8 and the reward probabilities of the actions at the blue state to a fixed value of 0.2, we obtain the results shown in Fig 2B instead of those shown in Fig 2A. (Similar results have already been observed by Smittenaar et al. [6] and Miller et al. [15].) Recall that these are computer-simulated model-free agents, who cannot use a model-based system to perform the task because they do not have one. Thus, this pattern cannot result from a shift between model-free and model-based influences on behavior.

The reason for this change is not that the reward probabilities are now fixed rather than variable. If we fix the reward probabilities to 0.5, we obtain the original pattern again, as shown in Fig 2C. In their original paper, Daw et al. [1] noted that the reward probabilities drift from trial to trial because this encourages participants to keep learning. Continued learning is a critical feature for testing many hypotheses, but it is not the feature that distinguishes model-free from model-based behavior.

The different model-free pattern in Fig 2B versus Fig 2A is caused by one final state being associated with a higher reward probability than the other. If actions taken at one final state are more often rewarded than actions taken at the other final state, the initial-state action that commonly leads to the most frequently rewarded final state will also be rewarded more often than the other initial-state action. This means that in trials that were rewarded after a common transition or unrewarded after a rare transition, corresponding to the outer bars of the plots, the agent usually chose the most rewarding initial-state action, and in trials that were rewarded after a rare transition or unrewarded after a common transition, corresponding to the inner bars of the plots, the agent usually chose the least rewarding initial-state action. Since one initial-state action is more rewarding than the other, model-free agents will learn to choose that action more often than the other, and thus, the stay probability for that action will be on average higher than the stay probability for the other action. This creates a tendency for the outer bars to be higher than the inner bars, and alters the pattern of model-free results relative to the canonical pattern by introducing an interaction between reward and transition. It does not alter the pattern of model-based results because model-based results already have higher outer bars and lower inner bars even if all reward probabilities are 0.5 (or stochastically drifting around 0.5).

Furthermore, unequal final-state reward probabilities will have an even greater effect on model-free agents with an eligibility trace parameter *λ* < 1 (Fig 2D). This is because the values of the initial-state actions are updated depending on the values of the final-state actions, which causes the action that takes the agent to the most rewarding final state to be updated to a higher value than the action that takes it to the least rewarding final state (see Equation 9 in the Methods section for details).

It also follows that if the reward probabilities of the final state-actions drift too slowly relative to the number of trials, model-free results will also exhibit an interaction between reward and transition. This is why the simulated results obtained by Miller et al. [15] using a simplified version of the two-step task do not exhibit the expected pattern; it is not because the task was simplified by only allowing one action at each final state. In that study, there was a 0.02 probability that the reward probabilities of the two final-state action (0.8 and 0.2) would be swapped, unless they had already been swapped in the previous 10 trials. If the swap probability is increased to 0.2 for a task with 250 trials, the canonical results are obtained instead (results not shown).

Despite changes in the expected pattern of model-free choices, it is still possible to use this modification of the task together with a logistic regression analysis to distinguish between model-free and model-based agents based on reward and transition. In order to do so, we simply need to include two more features in the analysis. As previously discussed, experimental data from two-stage tasks are typically analyzed by a logistic regression model, with *p*_stay_, the stay probability, as the dependent variable, and *x*_*r*_, a binary indicator of reward (+1 for rewarded, −1 for unrewarded), *x*_*t*_, a binary indicator of transition (+1 for common, −1 for rare), and *x*_*r*_*x*_*t*_, the interaction between reward and transition, as the independent variables:

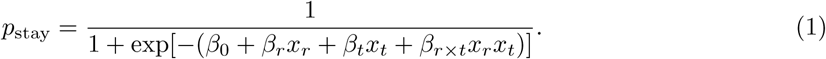

The levels of the independent variables were coded as +1 and −1 so that the meaning of the coefficients are easy to interpret: *β*_*r*_ indicates a main effect of reward, *β*_*t*_ indicates a main effect of transition, and *β*_*r×t*_ indicates an interaction between reward and transition. We applied this analysis to create all the plots presented so far, which can also be created from raw simulation data with similar results. In the modified task we just discussed, the *β*_*r×t*_ coefficient is positive for model-free agents, which does not allow us to distinguish between purely model-free and hybrid model-free/model-based agents.

We can, however, obtain an expected null *β*_*r×t*_ coefficient for purely model-free agents if we add two control variables to the analysis: *x*_*c*_, a binary indicator of the initial-state choice (+1 for left, −1 for right), and *x*_*f*_, a binary indicator of the final state (+1 for pink, −1 for blue):

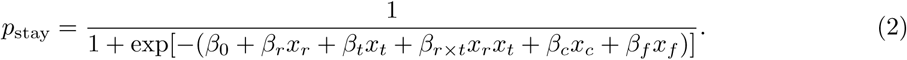

These two additional variables control for one initial-state choice having a higher stay probability than the other and for one final state having a higher reward probability than the other, respectively. The variable *x*_*f*_ is only necessary for model-free agents with *λ* < 1, because only in this case are the values of the initial-state actions updated depending on the values of the final-state actions.

By applying this extended logistic regression analysis to the same data used to generate Fig 2D and setting *x*_*c*_ = *x*_*f*_ = 0, we obtain Fig 2E, which is nearly identical to Fig 2A and Fig 2C. This result demonstrates that even though the original analysis fails to distinguish between model-free agents and hybrid agents, other analyses may succeed if they can extract more or different information from the data.

Another analysis that can be applied for this task is to fit a hybrid reinforcement learning model to the data and estimate the model-based weight (see [1] for details). A reinforcement learning model may be able to distinguish model-free and model-based behavior in this case without further modification. Kool et al. [18] describe another potential variation on the two-stage task in which model-free agents show interaction effects that are qualitatively similar to model-based agents, and those authors also suggest fitting reinforcement learning models to distinguish subtle differences between model-free and model-based behavior in such cases. However, we note that while reinforcement learning models will be more robust than logistic regression analyses in many cases, they will not be able to distinguish model-free and model-based actions equally well in every version of the two-stage task. Thus, computer simulation and parameter recovery exercises are advised when the data will be fit with reinforcement learning models as well.

### 2.2 Model-based agents will show main effects of transition in addition to transition by reward interactions under specific task conditions

When the final state probabilities do not sum to one, model-based agents will show both a main effect of transition and a transition by reward interaction. An example of these combined influences on model-based behavior can be seen in Fig 2F. This pattern was generated by modifying the original two-stage task so that the reward probability of all actions available at the pink and the blue states was 0.8. In this case, the reward probabilities of both final states are the same, and therefore, the stay probability of model-free agents is only affected by reward. On the other hand, the stay probability of model-based agents is not only affected by the interaction between reward and transition, but also by transition type itself. This main effect of transition can be seen in the right panel of Fig 2F by comparing the two outermost and innermost bars, which show that the common transitions (dark gray bars) lead to a lower stay probability relative to the corresponding rare transitions (light gray bars). This negative effect of common transitions on stay probabilities is because the sum of the reward probabilities of the final states, 0.8 and 0.8, is 1.6, which is greater than 1.

Fig 3 shows the relative extent to which the stay probabilities of model-based agents are influenced by transition type as a function of the sum of the reward probabilities at the final state. Let *p* be the value of the most valuable action at the pink state and *b* the value of the most valuable action at the blue state. The relative stay probabilities for model-based agents will be lower following common than rare transitions when *p* + *b* > 1. Conversely, relative stay probabilities for model-based agents will be higher following common than rare transitions when *p* + *b* < 1. Fig 3 shows the difference in stay probabilities between common and rare transitions as a function of both the sum of the final state reward probabilities and learning rate, *α*. Indeed, this graphic shows that model-based agents will show a main effect of transition in all cases except when *p* + *b* = 1. We explain the intuition and algebra behind this characteristic of our model-based agents in the following paragraphs.

**Fig 3:**
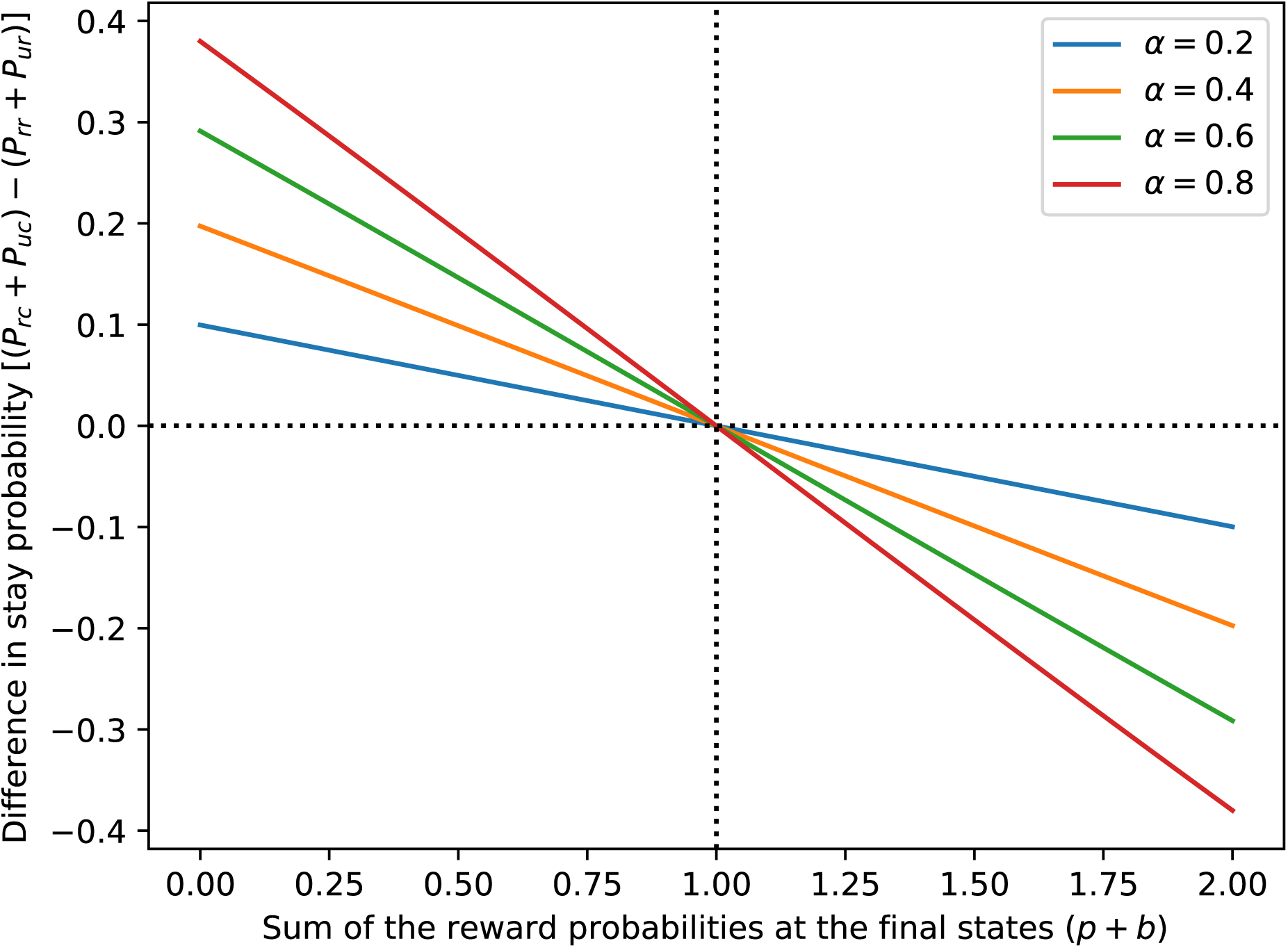
Difference in stay probability for model-based agents. Differences between the sum of the stay probabilities for model-based agents following common versus rare transitions (i.e., the sum of the dark gray bars minus the sum of the light gray bars) as a function of the sum of the reward probabilities at the final state (*p* + *b*). This specific example plot was generated assuming that final state reward probabilities are equal (*p* = *b*) and that the exploration-exploitation parameter in Equation 16 is *β* = 2.5. When computing the differences in stay probability on the y-axes, *P*_*rc*_ stands for the stay probability after a common transition and a reward, *P*_*uc*_ is the stay probability after a common transition and no reward, *P*_*rr*_ is the stay probability after a rare transition and a reward, and *P*_*ur*_ is the stay probability after a rare transition and no reward.

Model-based agents make initial-state decisions based on the difference, *p* − *b*, between the values of the most valuable actions available at the pink and blue states (this is a simplification; further details are given in the Methods section). As *p* − *b* increases, the agent becomes more likely to choose left, which commonly takes it to pink, and less likely to choose right, which commonly takes it to blue. This difference increases every time the agent experiences a common transition to pink and is rewarded (*p* increases) or experiences a rare transition to blue and is not rewarded (*b* decreases). Analogously, this difference decreases every time the agent experiences a common transition to blue and is rewarded (*b* increases) or experiences a rare transition to pink and is not rewarded (*p* decreases). This is why the model-based agent’s stay probabilities are affected by the interaction between reward and transition. But *p*−*b* may change *by different amounts* if the agent experiences a common transition and is rewarded versus if the agent experiences a rare transition and is not rewarded. If the agent experiences a common transition to pink and receives 1 reward, the difference between the final-state values changes from *p* − *b* to

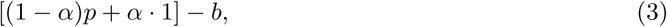

where 0 *≤ α ≤* 1 is the agent’s learning rate. If, on the other hand, the agent experiences a rare transition to blue and receives 0 rewards, the difference between the final-state values becomes

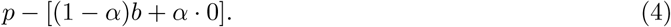

The two values are the same only if

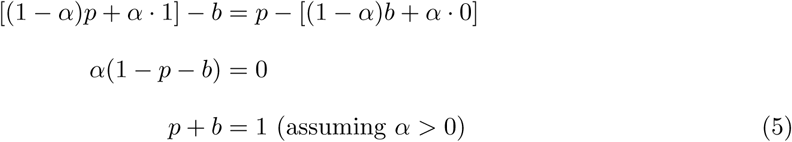

that is, when the sum of the final-state action values is 1. This is expected to occur when the actual reward probabilities of the final states sum to 1, as *p* and *b* estimate them. Thus, when the reward probabilities do not sum to 1, the outer bars of the stay probability plots may not be the same height. Similarly, *p* − *b* may change by different amounts if the agent experiences a common transition and is not rewarded versus if the agent experiences a rare transition and is rewarded, which also occurs when the reward probabilities do not sum to 1 (calculations not shown) and causes the inner bars of the stay probability plots to be different heights. In the S1 Appendix to this paper, we prove that this specifically creates a transition effect.

The end result is that the model-based behavior is not solely a function of the interaction between reward and transition, but also of the transition in many cases. Unlike our previous example, the main effect of transition cannot be corrected for by adding the initial-state choice and the final state as control variables. Fortunately, however, the original analysis can still be used to distinguish between model-free and model-based agents on this task because model-free agents exhibit only reward effects while model-based agents exhibit only transition and reward by transition interaction effects. According to Equations 29 and 29 in the Appendix, the transition coefficient *β*_*t*_ and the reward by transition interaction coefficient *β*_*r×t*_ of model-based agents are related so that *β*_*t*_ = (1 − *p* − *b*)*β*_*r×t*_. Therefore, if 1 ≠ *p* + *b*, both coefficients can be used to evaluate model-based control, since they are mathematically related by a known constant, which is determined by task design.

## 3 Discussion

The class of two-stage tasks pioneered by Daw et al. [1] has been instrumental in advancing efforts in the behavioral, computational, and biological sciences aimed at teasing apart the influences of model-free and model-based behavior and how the relative influences of these systems may change as a function of environmental context, biological development, and physical or mental health ([2, 3, 4, 5, 6, 7, 8, 9, 10, 11, 12, 13, 14, 15, 16, 17] among many others). The continued and expanded utilization of such tasks will require design modifications to better address specific new hypotheses and such efforts currently constitute an active and productive line of research across multiple scientific disciplines.

In the current paper, we have shown that slight modifications to established versions of the two-stage task design may deviate substantially from the expected patterns of results for both model-free and model-based agents when a logistic regression analysis is performed. Specifically, it is not a universal property of model-free and model-based learning that their stay probabilities are driven solely by rewards for model-free agents versus reward by transition interactions for model-based agents. Instead, the patterns of behavior produced by model-free and model-based agents are rather sensitive to changes in task features or learning algorithms. The two examples discussed here were just intended to illustrate this point, rather than present “flawed” versions of the two-stage paradigm to be avoided; indeed, it should be possible to use these modified tasks successfully in experiments, though it is important to keep in mind that they too rely on specific task features and parameterizations of the model-free and model-based learning algorithms.

Most importantly, there is a very straightforward means of avoiding potential design flaws or misinterpretations created by incorrect assumptions about the nature of model-free and model-based behavior in a given context—*test* how any changes in task design affect model-free and model-based agents’ choice patterns. Specifically, researchers who plan to use customized two-stage-style tasks in their work should always check by computer simulation of model-free and model-based agents what patterns each type of agent will produce in the new paradigm. It may be impossible to distinguish model-free from model-based choices with a logistic regression analysis containing only the previous outcome and transition as predictors. In this case, researchers can try adding additional relevant predictors to the analysis as we showed in Section 2.1. If a suitable set of logistic regression predictors cannot be found, it may be possible to analyze the data with a hybrid model-free/model-based reinforcement learning model.

It is obviously best to know if an extended logistic regression or reinforcement learning model can effectively achieve the analysis objectives from the outset, and thus, we recommend simulating and analyzing the behavior of model-based, model-free, or hybrid agents when planning to use a two-stage task. In order for *any* model to be able to distinguish between model-based and model-free behavior, it is necessary that the two algorithms make distinct choices in a sufficient number of trials. Such exercises in generating simulated data and analyzing them will allow researchers to tell if the data to be collected from a given task will contain enough information to allow retrieval of model parameters within the desired level of precision. More generally, they will allow researchers to better understand both the intended as well as potential unintended consequences of their design modifications *before* spending the time, effort, and money to acquire data from human participants or non-human animals. This will lead to better experimental designs that in turn yield more readily interpretable and informative conclusions about the question(s) of interest.

## 4 Methods

The code used to generate the results discussed in this paper is available at Github: https://github.com/carolfs/note_analysis_2stage_tasks

### 4.1 Task

The results were obtained by simulating model-free and model-based agents performing the two-stage task reported by Daw et al. [1] for 250 trials. In each trial, the agent first decides whether to perform the left or right action. Performing an action takes the agent to one of two final states, pink or blue. The left action takes the agent to pink with 0.7 probability (common transition) and to blue with 0.3 probability (rare transition). The right action takes the agent to blue with 0.7 probability (common transition) and to pink with 0.3 probability (rare transition). There are two actions available at final states. Each action has a different reward probability depending on whether the final state is pink or blue.

### 4.2 Simulation parameters

In the simulation of the two-stage task with drifting reward probabilities, all reward probabilities were initialized at a random value in the interval [0.25, 0.75] and drifted in each trial by the addition of random noise with distribution *𝒩* (*µ* = 0, *σ* = 0.025), with reflecting bounds at 0.25 and 0.75. Thus, the expected reward probability of final-state actions is 0.5. In the simulations of tasks with fixed reward probabilities, three different settings were used for the reward probabilities of the final-state actions: (1) 0.5 for all actions, (2) 0.8 for the actions available at the pink state and 0.2 for the actions available at the blue state, and (3) 0.8 for all actions.

The learning rate of the model-free agents was *α* = 0.5, the eligibility trace parameter was *λ* = 0.6 (for the case *λ* < 1) or *λ* = 1, and the exploration parameter was *β* = 5. The learning rate of model-based agents was *α* = 0.5 and the exploration parameter was *β* = 5. These parameter values are close to the median estimates in Daw et al. [1]. The values of all actions for all states were initialized at 0.

It should be noted, however, that all the explanations given for the observed results are based only on task design and mathematical calculations, not on the specific parameter values used in the simulations. Therefore, the study’s conclusions should not be affected by other parameters values, under the assumptions that agents are not making completely random choices (*β* > 0), that they learn from each outcome (*α* > 0) and retain this information in the long term (*α* < 1), and that the rewards obtained at the final states have a direct reinforcing effect on model-free choices at the initial state (*λ* > 0).

### 4.3 Model-free algorithm

Model-free agents were simulated using the SARSA(*λ*) algorithm [19, 1]. Specifically for two-stage tasks [1], the SARSA(*λ*) algorithm specifies that when an agent performs an initial-state action *a*_*i*_ at the initial state *s*_*i*_ (the index *i* stands for “initial”), then goes to the final state *s*_*f*_, performs the final-state action *a*_*f*_ (the index *f* stands for “final”) and receives a reward *r*, the model-free value *Q*_*MF*_ (*s*_*i*_, *a*_*i*_) of the initial-state action is updated as

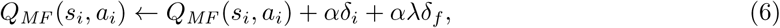

where

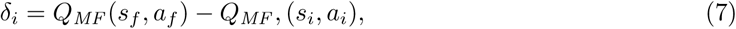

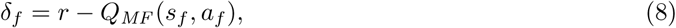

*α* is the learning rate and *λ* is the eligibility trace parameter [1]. Alternatively, the updating rule can be expressed in a single equation:

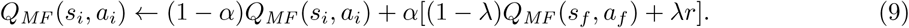

Since *λ* is a constant, this means that the value of an initial-state action is updated depending on the obtained reward and the value of the performed final-state action. If *λ* = 1, the equation becomes

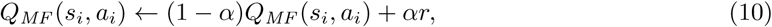

that is, the updated value depends only on the reward. The value *Q*_*MF*_ (*s*_*f*_, *a*_*f*_) of the final-state action is updated as

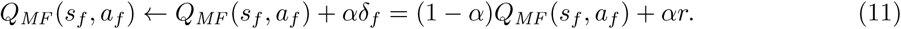

The probability *P* (*a*|*s*) that an agent will choose action *a* at state *s* is given by

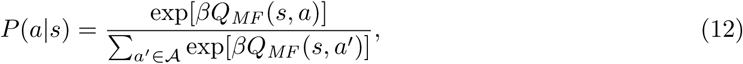

where 𝒜 is the set of all actions available at state *s* and *β* is an exploration-exploitation parameter [19].

### 4.4 Model-based algorithm

Model-based agents were simulated using the algorithm defined by Daw et al. [1]. Model-based agents make initial-state decisions based on the estimated value of the most valuable final-state actions and the transition probabilities. The value *Q*_*MB*_ (*s*_*i*_, *a*_*i*_) of an initial-state action *a*_*i*_ performed at the initial state *s*_*i*_ is

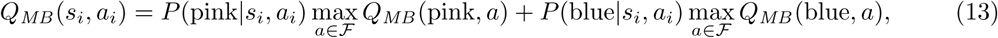

where *P* (*s*_*f*_ |*s*_*i*_, *a*_*i*_) is the probability of transitioning to the final state *s*_*f*_ by performing action *a*_*i*_ and ℱ is the set of actions available at the final states [1].

When the agent receives a reward, it will update the value of the final-state action *a*_*f*_ performed at state *s*_*f*_, *Q*_*MB*_ (*s*_*f*_, *a*_*f*_), according to the equation

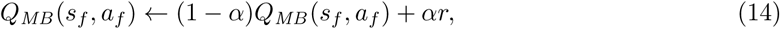

where *α* is the learning rate and *r* is the reward.

Let *p* = max_*a∈F*_ *Q*_*MB*_ (pink, *a*) and *b* = max_*a∈ℱ*_ *Q*_*MB*_ (blue, *a*). The probability *P* (left|*s*_*i*_) that the agent will choose the left action at the initial state *s*_*i*_ is given by

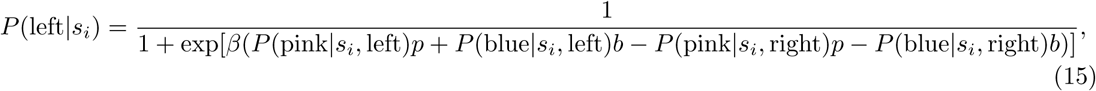

where *β* is an exploration-exploitation parameter. If each initial-state action transitions to a different final state with the same probability, e.g., *P*(pink|*s*_*i*_, left) = *P* (blue|*s*_*i*_, right) and hence *P*(pink|*s*_*i*_, right) = *P*(blue|*s*_*i*_, left), this equation is simplified to

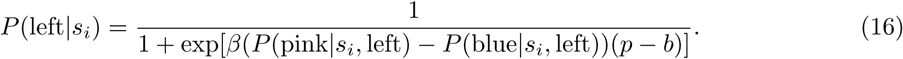

Hence, the agent’s probability of choosing left, the action that will take it more commonly to the pink state, increases with *p* − *b*.

### 4.5 Analysis

The simulation data were analyzed using the logistic regression models described in the Results section. 1,000 model-free and 1,000 model-based agents were simulated for each task modification discussed. The regression models were fitted to the data using the regularized logistic regression classifier with the liblinear algorithm from scikit-learn, a Python machine learning package [20].

## S1 Appendix

We will prove that if *p* + *b* ≠ 1, then there is a transition effect on the results of model-based agents. As explained in the Methods, if each initial-state action transitions to a different final state with the same probability, then the probability *P* (left|*s*_*i*_) of choosing left at the initial state *s*_*i*_ is given by

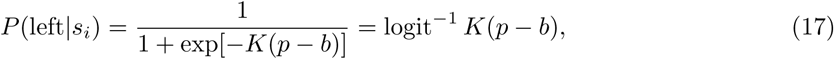

where *K* ≥ 0 is a constant that depends on the transition probabilities and the exploration-exploitation parameter.

According to the model-based reinforcement learning rule (Equation 14), if the agent chooses left, then experiences a common transition to pink and receives 1 reward, the stay probability *p*_stay_ (of choosing left again in the next trial) is given by

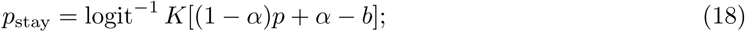

if instead the agent experiences a rare transition to blue and receives 1 reward, *p*_stay_ is given by

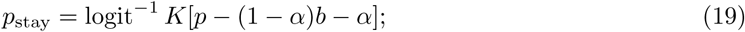

if the agent experiences a common transition to pink and receives 0 rewards, *p*_stay_ is given by

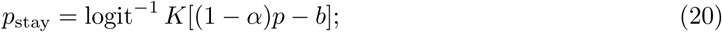

and if the agent experiences a rare transition to blue and receives 0 rewards, *p*_stay_ is given by

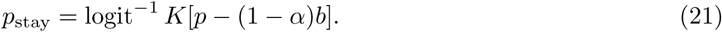

The logistic regression model, on the other hand, determines *p*_stay_ as a function *x*_*r*_ (*x*_*r*_ = +1 for 1 reward, *x*_*r*_ = −1 for 0 rewards in the previous trial) and *x*_*t*_ (*x*_*t*_ = +1 for a common transition, *x* = −1 for a rare transition in the previous trial):

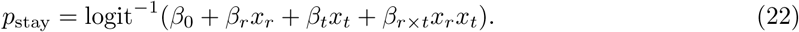

Since logit^−1^ is a one-to-one function, this implies that

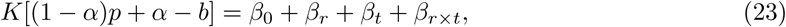

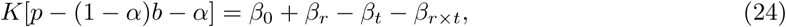

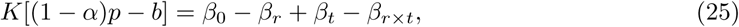

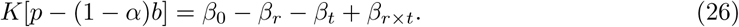

Solving this system for *β*_0_, *β*_*r*_, *β*_*t*_, and *β*_*r×t*_ yields

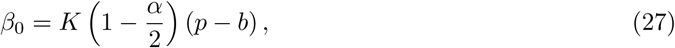

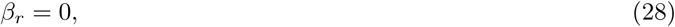

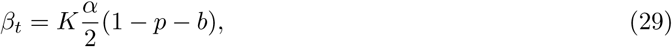

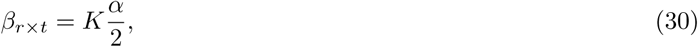

which implies that if *α* > 0, *K* > 0 and *p* + *b* ≠ 1, then *β*_*t*_ ≠ 0. This proof assumes that the agent chose left, but the same can be proved if the agent chose right, as in this example “left,” “right,” “pink,” and “blue” are arbitrary.

## References

[1] Nathaniel D. Daw, Samuel J. Gershman, Ben Seymour, Peter Dayan, and Raymond J. Dolan. Model-Based Influences on Humans’ Choices and Striatal Prediction Errors. Neuron, 69(6): 1204–1215, mar 2011. ISSN 08966273. doi: 10.1016/j.neuron.2011.02.027. URL http://www.cell.com/neuron/abstract/S0896-6273(11)00125-5http://www.pubmedcentral.nih.gov/articlerender.fcgi?artid=3077926{&}tool=pmcentrez{&}rendertype=abstracthttp://linkinghub.elsevier.com/retrieve/pii/S0896627311001255.

[2] Klaus Wunderlich, Peter Smittenaar, and Raymond J. Dolan. Dopamine Enhances Model-Based over Model-Free Choice Behavior. Neuron, 75(3):418–424, aug 2012. ISSN 08966273. doi: 10.1016/j.neuron.2012.03.042. URL http://linkinghub.elsevier.com/retrieve/pii/S0896627312005272.

[3] Ben Eppinger, Maik Walter, Hauke R. Heekeren, and Shu-Chen Li. Of goals and habits: age-related and individual differences in goal-directed decision-making. Frontiers in Neuroscience, 7, 2013. ISSN 1662-453X. doi: 10.3389/fnins.2013.00253. URL http://journal.frontiersin.org/article/10.3389/fnins.2013.00253/abstract.

[4] A. R. Otto, C. M. Raio, A. Chiang, E. A. Phelps, and N. D. Daw. Working-memory capacity protects model-based learning from stress. Proceedings of the National Academy of Sciences, 110(52):20941–20946, dec 2013. ISSN 0027-8424. doi: 10.1073/pnas.1312011110. URL http://www.pnas.org/cgi/doi/10.1073/pnas.1312011110.

[5] A. Ross Otto, Samuel J. Gershman, Arthur B. Markman, and Nathaniel D. Daw. The Curse of Planning. Psychological Science, 24(5):751–761, may 2013. ISSN 0956-7976. doi: 10.1177/0956797612463080. URL http://journals.sagepub.com/doi/10.1177/0956797612463080.

[6] Peter Smittenaar, Thomas H.B. FitzGerald, Vincenzo Romei, Nicholas D. Wright, and Raymond J. Dolan. Disruption of Dorsolateral Prefrontal Cortex Decreases Model-Based in Favor of Model-free Control in Humans. Neuron, 80(4):914–919, nov 2013. ISSN 08966273. doi: 10.1016/j.neuron.2013.08.009. URL http://linkinghub.elsevier.com/retrieve/pii/S0896627313007204.

[7] Amir Dezfouli and Bernard W. Balleine. Actions, Action Sequences and Habits: Evidence That Goal-Directed and Habitual Action Control Are Hierarchically Organized. PLoS Computational Biology, 9(12):e1003364, dec 2013. ISSN 1553-7358. doi: 10.1371/journal.pcbi.1003364. URL http://dx.plos.org/10.1371/journal.pcbi.1003364.

[8] Miriam Sebold, Lorenz Deserno, Stefan Nebe, Daniel J. Schad, Maria Garbusow, Claudia Hägele, Jürgen Keller, Elisabeth Jünger, Norbert Kathmann, Michael Smolka, Michael A. Rapp, Florian Schlagenhauf, Andreas Heinz, and Quentin J.M. Huys. Model-Based and Model-Free Decisions in Alcohol Dependence. Neuropsychobiology, 70(2):122–131, 2014. ISSN 0302-282X. doi: 10.1159/000362840. URL https://www.karger.com/?doi=10.1159/000362840.

[9] V Voon, K Derbyshire, C Rück, M A Irvine, Y Worbe, J Enander, L R N Schreiber, C Gillan, N A Fineberg, B J Sahakian, T W Robbins, N A Harrison, J Wood, N D Daw, P Dayan, J E Grant, and E T Bullmore. Disorders of compulsivity: a common bias towards learning habits. Molecular Psychiatry, 20(3):345–352, mar 2015. ISSN 1359-4184. doi: 10.1038/mp.2014.44. URL http://www.nature.com/doifinder/10.1038/mp.2014.44.

[10] Bradley B Doll, Katherine D Duncan, Dylan A Simon, Daphna Shohamy, and Nathaniel D Daw. Model-based choices involve prospective neural activity. Nature Neuroscience, 18(5):767–772, mar 2015. ISSN 1097-6256. doi: 10.1038/nn.3981. URL http://www.nature.com/doifinder/10.1038/nn.3981.

[11] Fiery Cushman and Adam Morris. Habitual control of goal selection in humans. Proceedings of the National Academy of Sciences, 112(45):13817–13822, nov 2015. ISSN 0027-8424. doi: 10.1073/pnas.1506367112. URL http://www.pnas.org/lookup/doi/10.1073/pnas.1506367112.

[12] A. Ross Otto, Anya Skatova, Seth Madlon-Kay, and Nathaniel D. Daw. Cognitive Control Predicts Use of Model-based Reinforcement Learning. Journal of Cognitive Neuroscience, 27(2):319–333, feb 2015. ISSN 0898-929X. doi: 10.1162/jocn_a_00709. URL www.mitpressjournals.org/doi/abs/10.1162/jocn{_}a{_}00709 http://www.mitpressjournals.org/doi/10.1162/jocn{_}a{_}00709.

[13] Lorenz Deserno, Quentin J. M. Huys, Rebecca Boehme, Ralph Buchert, Hans-Jochen Heinze, Anthony A. Grace, Raymond J. Dolan, Andreas Heinz, and Florian Schlagenhauf. Ventral striatal dopamine reflects behavioral and neural signatures of model-based control during sequential decision making. Proceedings of the National Academy of Sciences, 112(5):1595–1600, feb 2015. ISSN 0027-8424. doi: 10.1073/pnas.1417219112. URL http://www.pnas.org/lookup/doi/101073/pnas.1417219112.

[14] Claire M. Gillan, A. Ross Otto, Elizabeth A. Phelps, and Nathaniel D. Daw. Model-based learning protects against forming habits. Cognitive, Affective, & Behavioral Neuroscience, 15(3):523–536, sep 2015. ISSN 1530-7026. doi: 10.3758/s13415-015-0347-6. URL http://link.springer.com/10.3758/s13415-015-0347-6 http://link.springer.com/article/10.3758/s13415-015-0347-6{%}5Cn http://link.springer.com/article/10.3758{%}2Fs13415-015-0347-6{%}5Cn http://link.springer.com/content/pdf/10.3758{%}2Fs13415-015-0347-6.pdf.

[15] Kevin J Miller, Carlos D Brody, and Matthew M Botvinick. Identifying Model-Based and Model-Free Patterns in Behavior on Multi-Step Tasks. bioRxiv, page 14, 2016. doi: 10.1101/096339. URL https://doi.org/10.1101/096339.

[16] Wouter Kool, Samuel J. Gershman, and Fiery A. Cushman. Cost-Benefit Arbitration Between Multiple Reinforcement-Learning Systems. Psychological Science, page 095679761770828, jul 2017. ISSN 0956-7976. doi: 10.1177/0956797617708288. URL http://journals.sagepub.com/doi/10.1177/0956797617708288.

[17] Laurel S. Morris, Kwangyeol Baek, and Valerie Voon. Distinct cortico-striatal connections with subthalamic nucleus underlie facets of compulsivity. Cortex, 88:143–150, mar 2017. ISSN 00109452. doi: 10.1016/j.cortex.2016.12.018. URL http://linkinghub.elsevier.com/retrieve/pii/S0010945216303719.

[18] Wouter Kool, Fiery A. Cushman, and Samuel J. Gershman. When Does Model-Based Control Pay Off? PLOS Computational Biology, 12(8):e1005090, aug 2016. ISSN 1553-7358. doi: 10.1371/journal.pcbi.1005090. URL http://dx.plos.org/10.1371/journal.pcbi.1005090.

[19] Richard S. Sutton and Andrew G. Barto. Reinforcement Learning: An Introduction. A Bradford Book, first edition, 1998.

[20] F Pedregosa, G Varoquaux, A Gramfort, V Michel, B Thirion, O Grisel, M Blondel, P Prettenhofer, R Weiss, V Dubourg, J Vanderplas, A Passos, D Cournapeau, M Brucher, M Perrot, and E Duchesnay. Scikit-learn: Machine Learning in *{*P*}*ython. Journal of Machine Learning Research, 12:2825–2830, 2011.

